# CERMEP-IDB-MRXFDG: A database of 37 normal adult human brain [^18^F]FDG PET, T1 and FLAIR MRI, and CT images available for research

**DOI:** 10.1101/2020.12.15.422636

**Authors:** Inés Mérida, Julien Jung, Sandrine Bouvard, Didier Le Bars, Sophie Lancelot, Franck Lavenne, Caroline Bouillot, Jérôme Redouté, Alexander Hammers, Nicolas Costes

## Abstract

We present a database of cerebral PET FDG and anatomical MRI for 37 normal adult human subjects (CERMEP-IDB-MRXFDG).

Thirty-nine participants underwent [^18^F]FDG PET/CT and MRI, resulting in [^18^F]FDG PET, T1 MPRAGE MRI, FLAIR MRI, and CT images. Two participants were excluded after visual quality control. We describe the acquisition parameters, the image processing pipeline and provide participants’ individual demographics (mean age 38 ± 11.5 years, range 23-65, 20 women). Volumetric analysis of the 37 T1 MRIs showed results in line with the literature. A leave-one-out assessment of the 37 FDG images using Statistical Parametric Mapping (SPM) yielded a low number of false positives after exclusion of artefacts.

The database is stored in three different formats, following the BIDS common specification: 1) DICOM (data not processed), 2) NIFTI (multimodal images coregistered to PET subject space), 3) NIFTI normalized (images normalized to MNI space).

*Bona fide* researchers can request access to the database via a short form.

## Introduction

Imaging databases are very useful to re-analyse data in a different context, to increase the number of subjects of a study, and to develop new methods. Imaging databases play a crucial role in numerous analysis methods that rely in the comparison between the data of a group or of an individual and a group of reference. This includes studies using a normative database for analysis and quantification purposes, machine learning approaches, multi-atlas techniques, and validation of image processing pipelines. Databases with different modalities per participant also allow approaches that derive “missing” modalities, e.g. creating pseudo-CTs for attenuation correction in PET-MR (Merida et al. 2017; Burgos et al. 2014; Yaakub, McGinnity, Beck, et al. 2019; Ladefoged et al. 2019).

In the last years, an increasing number of neuroimaging databases has been made available. These databases generally consist of MR images (such as ADNI http://adni.loni.usc.edu, OASIS https://www.oasis-brains.org; for a review see (Coupé et al. 2017)). There is also a large database of PET from the Copenhagen group, CIMBI, containing mainly serotonine receptor PET and associated data (Knudsen et al. 2016). We are aware of very few datasets for [^18^F]fluorodeoxyglucose ([^18^F]FDG) PET imaging that have been published (Wei et al. 2018) or are available on request (Archambaud et al. 2013; Eusebio et al. 2012; Alzheimer’s Disease Neuroimaging Initiative, ADNI http://adni.loni.usc.edu).

Acquisition of imaging data, such as MRI scanning and in particular PET imaging that requires the injection of a radiotracer, represents an important logistical and monetary cost. In addition, participants have to consent to data acquisition and dissemination, and many countries have restrictions on using ionising radiation in healthy controls, adding to difficulties in acquiring such databases. Database sharing thus contributes to reduce research costs and reduces radiation exposure of healthy controls.

In order to make database sharing more efficient, the scientific community has implemented a database standardisation to organize and describe the data (Brain Imaging Data Structure (BIDS), https://bids.neuroimaging.io, (Gorgolewski et al. 2016)). In this work we introduce a multi-modal database of 37 healthy subjects constructed with MRI, CT and [^18^F]FDG PET images to BIDS standard. We have obtained ethical permission to share the data on request.

## Materials & Methods

### Recruitment and cohort characteristics

All enrolled subjects provided written informed consent to participate in the study (EudraCT: 2014-000610-56). The subjects were informed that their anonymized images could be used for methodological development and had been given the option to oppose this use of their data. The inclusion criteria were adult healthy subject and aged between 20 and 65 years. Exclusion criteria were (1) children and adults older than 65 years, (2) woman of childbearing potential without effective contraception, (3) history of neurological disorders, (4) any contraindication for MRI scanning, (5) active infectious disease. Thirty-nine subjects were included in the study. Each subject had a T1-weighted MRI, a T2 fluid-attenuated inversion recovery (FLAIR) MRI and an [^18^F]FDG PET/CT brain scan. The subjects’ MR and PET images were visually reviewed by two neurologists for conspicuous brain abnormalities. Two subjects showing brain lesions on the MR images (one probable insular cavernoma, one cerebellar lesion with hyperintense signal in the FLAIR sequence suggesting possible inflammatory disease of the central nervous system) were excluded from the database.

### MRI acquisition and reconstruction

MRI sequences were obtained on a Siemens Sonata 1.5 T scanner. Three-dimensional anatomical T1-weighted sequences (MPRAGE) were acquired in sagittal orientation (TR 2400 ms, TE 3.55 ms, inversion time 1000 ms, flip angle 8°). The images were reconstructed into a 160 x 192 x 192 matrix with voxel dimensions of 1.2 x 1.2 x 1.2 mm^3^ (axial field of view 230.4 mm). Sagittal Fluid-Attenuated Inversion Recovery (FLAIR, Hajnal et al. 1993) images (TR 6000 ms, TE 354 ms, Inversion time 2200 ms, flip angle 180°) were acquired with a 176 x 196 x 256 matrix and a voxel size of 1.2 x 1.2 x 1.2 mm^3^ (axial field of view 307.2 mm).

### PET and CT acquisition and reconstruction

PET and CT data were acquired on a Siemens Biograph mCT64. During the uptake period, participants were instructed to rest with their eyes closed and without auditory stimulation. PET data acquisition started 50 min after the injection of 122.30 ± 21.29 MBq of [^18^F]FDG (individual doses are provided in the demographics table) and lasted 10 min. PET images were reconstructed using 3D ordinary Poisson-ordered subsets expectation maximization (OP-OSEM 3D), incorporating the system point spread function and time of flight, and using 12 iterations and 21 subsets (Siemens’ “HD reconstruction”). Data correction (normalization, attenuation and scatter correction) was fully integrated within the reconstruction process. Gaussian post-reconstruction 3D filtering (FWHM = 4 mm isotropic) was applied to all PET images. Reconstructions were performed with a zoom of 2 yielding a voxel size of 2.04 x 2.04 x 2.03 mm^3^ in a matrix of 200 x 200 x 109 voxels (axial field of view 221.27 mm). Low-dose CT images for attenuation correction were acquired with a tube voltage of 100 keV and reconstructed in a 512×512×233 matrix with a voxel size of 0.6 x 0.6 x 1.5 mm^3^ (axial field of view 349.5 mm).

### Processing pipeline

#### Data anonymisation and pre-processing

Data anonymisation was performed on the DICOM files using the *gdcmanon* function (http://gdcm.sourceforge.net/html/gdcmanon.html). DICOM files were converted to NIFTI format with *dcm2niix* software (https://github.com/rordenlab/dcm2nii).

The background of CT images was cleaned in order to remove the scanner table and other objects such as the pillow included in the background of the image. For this, a binary mask of the head of the subject was automatically generated following a procedure described in (Merida et al 2015) using tools from the FSL (Version 6.0, https://fsl.fmrib.ox.ac.uk/fsl/fslwiki/) and NiftySeg (http://cmictig.cs.ucl.ac.uk/wiki/index.php/NiftySeg) suites. Finally, the binary mask was applied to the CT image.

#### Coregistration

As first step, the origin of each NIFTI image was set to the matrix centre. Then, CT, T1 MRI and FLAIR MRI images were coregistered to the [^18^F]FDG PET image using the *Coregister & Estimate* function from the SPM 12 toolbox (https://www.fil.ion.ucl.ac.uk/spm/software/spm12/).

#### Spatial normalisation

All images were normalized to MNI space through the tissue classification into grey and white matter probability maps of the T1 image. For that, individual subject’s deformation fields were calculated by the *Segment* function of SPM 12,(Ashburner and Friston 2005) from the T1 images previously coregistered to the PET image (but not resliced to preserve native resolution). Transformations for MR to PET space coregistration and PET to MNI space normalisation were concatenated and applied at once to avoid an intermediate resampling of the MRI data. All normalized images were resampled at 1×1×1 mm using 4^th^ degree B-spline interpolation.

#### Intensity normalisation

Reconstructed PET images were normalized by the subjects’ weight and injected dose to obtain Standard Uptake Value (SUV) images (radioactivity concentration [kBq/cm^3^] / (dose [kBq] / weight [kg])). In addition, reconstructed PET images were normalized by each subject’s mean activity within the intracranial volume (ICV) mask provided by SPM12 to obtain Standard Uptake Value ratio (SUVr).

### Regional analysis

The T1 MR images were anatomically segmented into 83 regions using the Hammers_mith maximum probability atlas n30r83, which is based on the mutli-atlas fusion of 30 manually delineated MRIs of healthy young adults (Hammers et al. 2003; Gousias et al. 2008), available at http://brain-development.org. The atlas was wrapped to each individual MRI space *via* the inverse transformation of the deformation fields from subject’s space to the MNI space computed at the spatial normalisation step. Grey matter and white matter probability maps obtained with the *Segment* function were thresholded at 0.5 and combined with the 83-ROI anatomical segmentation in order to separate their grey and white matter parts, expect for pure white matter regions like the corpus callosum, and pure grey matter regions like the basal ganglia.

Mean regional SUV and SUVr were extracted in a selection of grey matter anatomical regions of the Hammers_mith segmentation.

### Leave-one-out SPM analysis on [^18^F]FDG images

Leave-one-out ANCOVA was performed on SPM12 in order to compare each subject (healthy control) of the database to the others.

For the statistical analysis, PET images were smoothed with a Gaussian filter at 8mm FWHM. We used age and the global mean calculated within the intracranial volume mask as covariates. Two different contrasts were explored: Hyper-metabolism, i.e. activity of one subject > activity of the remaining subjects in the database, and hypo-metabolism, i.e. activity of one subject < activity of the remaining subjects in the database. Significant differences where defined at p < 0.05 FWE at the cluster level.

The database outliers were assessed with three criteria, for both hypometabolism and hypermetabolism.

– Subject-level: number of subjects with significant differences / total number of subjects in the database x 100
– Cluster-level: total number of significant clusters across all subjects / average number of resolution elements (resells) in the mask x 100
– Voxel-level: total number of voxels among the significant clusters across all subjects / number of voxels in the SPM mask x 100

## Results

### Database IDB-MRXFDG

The final database consists of 37 participants (17 male / 20 female, mean age ± SD, 38.11 ± 11.36 years; range, 23-65 years). Each participant has [^18^F]FDG PET, T1 MRI, FLAIR MRI, and CT images. An example of coregistered T1, FLAIR, CT and [^18^F]FDG PET images in the subject space are shown in Figure 1 and the same images in normalized space are shown in Figure 2.

**Figure 1:**
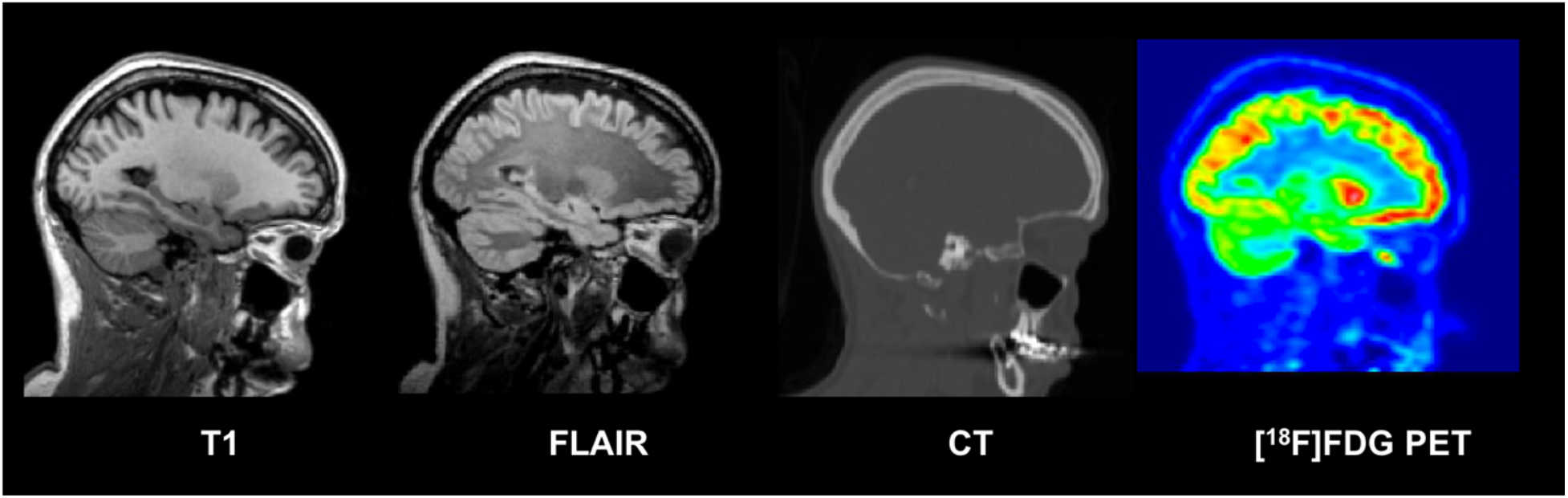
Example of coregistered T1 MRI, FLAIR MRI, CT and [^18^F]FDG PET images (sagittal plane) for one subject of the database

**Figure 2:**
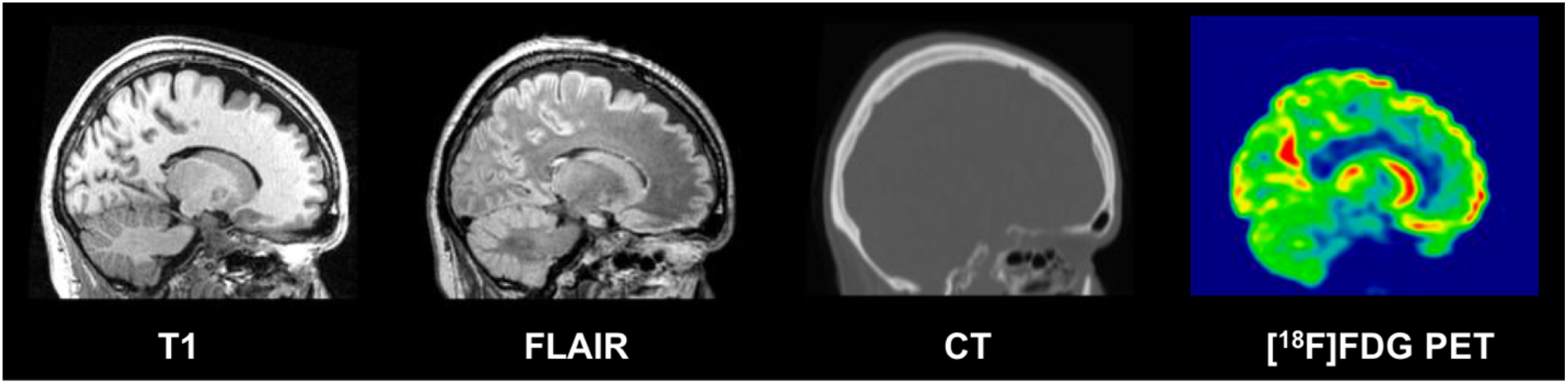
Example of normalized T1 MRI, FLAIR MRI, CT and [^18^F]FDG PET images (sagittal plane) in MNI space, for one subject of the database

Table 1 summarizes the demographic information for each participant: subject ID, acquisition date, age of the participant at the time of the imaging session, sex, weight, size, injected dose of [^18^F]FDG, handedness and a comment if any hypersignal was observed on the FLAIR MRI.

**Table 1:** Demographics table

| Subject ID | Birth date (year) | Age | Sex | Weight (kg) | Size (cm) | Injected Dose (MBq) | Mean activity in ICV mask (MBq) | Handedness |
| --- | --- | --- | --- | --- | --- | --- | --- | --- |
| sub-0001 | 1980 | 35 | F | 81 | 163 | 149 | 6911 | R |
| sub-0002 | 1957 | 58 | F | 66 | 160 | 102 | 5781 | R |
| sub-0003 | 1979 | 36 | M | 88 | 174 | 144 | 5980 | R |
| sub-0004 | 1963 | 51 | M | 72 | - | 124 | 6551 | R |
| sub-0005 | 1981 | 33 | M | 110 | 180 | 147 | 4925 | R |
| sub-0006 | 1974 | 41 | M | 76 | 176 | 119 | 5590 | R |
| sub-0007 | 1975 | 40 | F | 60 | 170 | 118 | 8271 | R |
| sub-0008 | 1988 | 27 | F | 53 | - | 95 | 6889 | R |
| sub-0009 | 1986 | 28 | F | 66 | - | 135 | 7974 | R |
| sub-0010 | 1971 | 43 | M | 117 | 170 | 168 | 6137 | R |
| sub-0011 | 1988 | 27 | F | 53 | 161 | 99 | 6944 | R |
| sub-0012 | 1990 | 24 | F | 63 | 168 | 123 | 7911 | R |
| sub-0013 | 1988 | 27 | M | 74 | 178 | 135 | 7798 | R |
| sub-0014 (*) | 1964 | 50 | F | 55 | 157 | 94 | 2574 | R |
| sub-0015 | 1981 | 34 | M | 62 | 170 | 109 | 6032 | R |
| sub-0016 (*) | 1949 | 65 | F | 70 | 170 | 108 | 4849 | R |
| sub-0017 | 1958 | 56 | M | 89 | 185 | 140 | 5539 | L |
| sub-0018 | 1969 | 45 | M | 71 | 168 | 132 | 4444 | R |
| sub-0019 | 1989 | 25 | F | 61 | 161 | 109 | 7755 | R |
| sub-0020 | 1973 | 42 | M | 72 | 185 | 135 | 6966 | L |
| sub-0021 | 1990 | 25 | M | 63 | 178 | 110 | 3119 | R |
| sub-0022 | 1990 | 25 | M | 72 | 178 | 133 | 7075 | R |
| sub-0023 | 1974 | 41 | F | 54 | 168 | 99 | 5970 | R |
| sub-0024 | 1976 | 39 | M | 118 | 199 | 167 | 6528 | R |
| sub-0025 (*) | 1967 | 48 | F | 83 | 168 | 149 | 5811 | R |
| sub-0026 | 1988 | 27 | F | 70 | - | 134 | 7656 | R |
| sub-0027 | 1963 | 52 | F | 57 | - | 107 | 6647 | R |
| sub-0028 | 1969 | 46 | F | 53 | 173 | 95 | 5481 | L |
| sub-0029 | 1991 | 23 | M | 80 | 178 | 136 | 7824 | R |
| sub-0030 | 1989 | 25 | M | 72 | 175 | 124 | 7384 | R |
| sub-0031 | 1979 | 35 | F | 58 | - | 112 | 6621 | R |
| sub-0032 | 1981 | 34 | M | 80 | 178 | 136 | 7011 | R |
| sub-0033 | 1955 | 60 | F | 48 | 159 | 95 | 6151 | R |
| sub-0034 | 1984 | 31 | F | 58 | 166 | 106 | 6119 | L |
| sub-0035 | 1969 | 46 | F | 48 | 167 | 96 | 6066 | R |
| sub-0036 | 1982 | 33 | M | 79 | 180 | 149 | 7253 | R |
| sub-0037 | 1981 | 33 | F | 47 | 163 | 92 | 7355 | R |
| **Mean** |  | 38.11 |  | 70.24 | 171.81 | 122 | 6375 |  |
| **SD** |  | 11.49 |  | 17.37 | 9.11 | 21 | 1276 |  |
| **Min** |  | 23 |  | 48 | 157 | 94 | 2574 |  |
| **Max** |  | 65 |  | 118 | 199 | 168 | 8271 |  |

sub-0014 : Hyperintense FLAIR signals in white matter (corona radiata) suggesting benign age related white matter hyperintensities (WMHs)
sub-0016 : Hyperintense FLAIR signals in white matter (corona radiata and frontal subcortical structures) suggesting benign age related white matter hyperintensities
sub-0025 : Hyperintense FLAIR signals in white matter (corona radiata and frontal subcortical structures) suggesting benign age related white matter hyperintensities
For those 3 subjects, the location and MRI changes observed in FLAIR sequences are typical findings of WMHs with diffuse areas of high signal intensity (hence, “hyperintense”) on T2-weighted or FLAIR sequences. Those WMHs are typically interpreted as a surrogate of cerebral small vessel disease. Due to the high prevalence of those MRI changes in asymptomatic subjects above 50 years, the PET images of those subjects were included in the database.

The database is available in three different formats, following the BIDS common specification:

– DICOM (data not processed)
– NIFTI (multimodal images coregistered to PET subject space)
– NIFTI normalized (images normalized to MNI space)

Table 2 lists the regional volumes obtained via the Hammers_mith maximum probability atlas. Coefficients of variation were as expected, without obvious outliers. The structure sizes were also in line with expectations (Hammers et al. 2003; Gousias et al. 2008).

**Table 2:** Regional volumes in native space (in cm^3^). Each paired region is composed of left and right sub-regions. The short names are expanded in Table A3 in the Appendix.

| Structure name | Grey matter | White matter | Structure name | Grey matter | White matter |
| --- | --- | --- | --- | --- | --- |
| | Mean ± SD | Mean ± SD | | Mean ± SD | Mean ± SD |
| **Temporal Lobe** |  |  | **Occipital Lobe** |  |  |
| Hippocampus_r | 2.37 ± 0.28 | - | OL_rest_lat_l | 21.95 ± 2.40 | 18.27 ± 2.83 |
| Hippocampus_l | 2.11 ± 0.26 | - | OL_rest_lat_r | 22.42 ± 2.54 | 19.36 ± 3.19 |
| Amygdala_r | 1.43 ± 0.17 | - | OL_ling_G_l | 7.56 ± 0.93 | 3.94 ± 0.70 |
| Amygdala_l | 1.52 ± 0.18 | - | OL_ling_G_r | 8.16 ± 0.91 | 3.97 ± 0.79 |
| Ant_TL_med_r | 5.48 ± 0.64 | 1.24 ± 0.28 | OL_cuneus_l | 5.31 ± 0.66 | 3.34 ± 0.61 |
| Ant_TL_med_l | 5.27 ± 0.61 | 1.22 ± 0.26 | OL_cuneus_r | 5.72 ± 0.70 | 3.09 ± 0.67 |
| Ant_TL_inf_lat_r | 2.77 ± 0.39 | 0.55 ± 0.19 | **Parietal Lobe** | | |
| Ant_TL_inf_lat_l | 2.58 ± 0.41 | 0.49 ± 0.17 | PL_rest_l | 21.37 ± 2.52 | 16.01 ± 2.40 |
| G_paraH_amb_r | 3.15 ± 0.37 | 0.98 ± 0.13 | PL_rest_r | 21.25 ± 2.48 | 15.95 ± 2.28 |
| G_paraH_amb_l | 3.27 ± 0.42 | 0.99 ± 0.18 | PL_postce_G_l | 11.69 ± 1.58 | 14.69 ± 1.96 |
| G_sup_temp_cent_r | 7.62 ± 0.93 | 5.33 ± 0.79 | PL_postce_G_r | 10.80 ± 1.43 | 13.82 ± 1.76 |
| G_sup_temp_cent_l | 7.64 ± 0.96 | 5.26 ± 0.80 | PL_sup_pa_G_l | 19.50 ± 2.10 | 17.65 ± 2.75 |
| G_tem_midin_r | 11.24 ± 1.33 | 5.95 ± 0.95 | PL_sup_pa_G_r | 20.51 ± 2.15 | 18.42 ± 3.05 |
| G_tem_midin_l | 10.78 ± 1.30 | 5.87 ± 1.03 | **Central Structures** | | |
| G_occtem_la_r | 3.42 ± 0.39 | 0.92 ± 0.24 | CaudateNucl_l | 4.24 ± 0.52 | - |
| G_occtem_la_l | 3.41 ± 0.41 | 0.94 ± 0.24 | CaudateNucl_r | 4.33 ± 0.54 | - |
| PosteriorTL_l | 26.41 ± 3.01 | 17.13 ± 2.44 | NuclAccumb_l | 0.36 ± 0.05 | - |
| PosteriorTL_r | 27.52 ± 3.05 | 17.48 ± 2.39 | NuclAccumb_r | 0.30 ± 0.04 | - |
| G_sup_temp_ant_l | 3.28 ± 0.42 | 0.67 ± 0.19 | Putamen_l | 4.92 ± 0.57 | - |
| G_sup_temp_ant_r | 3.19 ± 0.39 | 0.62 ± 0.17 | Putamen_r | 4.76 ± 0.54 | - |
| **Posterior Fossa** | | | Thalamus_l | 7.41 ± 0.87 | - |
| Cerebellum_r | 44.49 ± 5.10 | 12.34 ± 1.63 | Thalamus_r | 7.25 ± 0.86 | - |
| Cerebellum_l | 44.36 ± 5.13 | 12.39 ± 1.63 | Pallidum_l | 1.30 ± 0.16 | - |
| Brainstem | 23.05 ± 2.82 | - | Pallidum_r | 1.31 ± 0.16 | - |
| **Frontal Lobe** | | | Corp_Callosum | 20.83 ± 2.54 | - |
| FL_mid_fr_G_l | 23.21 ± 3.01 | 24.36 ± 3.67 | S_nigra_l | 0.32 ± 0.04 | - |
| FL_mid_fr_G_r | 23.59 ± 2.95 | 24.52 ± 3.57 | S_nigra_r | 0.32 ± 0.04 | - |
| FL_precen_G_l | 13.11 ± 1.76 | 18.47 ± 2.32 | **Ventricles** | | |
| FL_precen_G_r | 13.04 ± 1.84 | 18.55 ± 2.37 | BodyVentricle_r | 7.31 ± 1.01 | - |
| FL_OFC_AOG_l | 3.87 ± 0.51 | 1.45 ± 0.39 | BodyVentricle_l | 7.98 ± 1.01 | - |
| FL_OFC_AOG_r | 3.89 ± 0.51 | 1.45 ± 0.35 | TemporaHorn_r | 0.63 ± 0.08 | - |
| FL_inf_fr_G_l | 10.63 ± 1.34 | 5.96 ± 1.12 | TemporaHorn_l | 0.49 ± 0.06 | - |
| FL_inf_fr_G_r | 10.15 ± 1.24 | 5.50 ± 1.05 | ThirdVentricl | 0.94 ± 0.13 | - |
| FL_sup_fr_G_l | 27.15 ± 3.58 | 20.00 ± 3.10 | **Insula and Cingulate gyri** | | |
| FL_sup_fr_G_r | 27.33 ± 3.63 | 19.71 ± 3.12 | Insula_l | 15.42 ± 1.77 | - |
| FL_OFC_MOG_l | 3.95 ± 0.53 | 1.83 ± 0.37 | Insula_r | 15.37 ± 1.79 | - |
| FL_OFC_MOG_r | 3.93 ± 0.48 | 1.63 ± 0.30 | G_cing_ant_sup_l | 5.56 ± 0.74 | 1.66 ± 0.43 |
| FL_OFC_LOG_l | 2.13 ± 0.31 | 0.92 ± 0.22 | G_cing_ant_sup_r | 5.29 ± 0.69 | 1.67 ± 0.40 |
| FL_OFC_LOG_r | 2.39 ± 0.37 | 0.91 ± 0.24 | G_cing_post_l | 5.07 ± 0.67 | 2.52 ± 0.46 |
| FL_OFC_POG_l | 3.14 ± 0.41 | 1.34 ± 0.28 | G_cing_post_r | 4.95 ± 0.60 | 2.36 ± 0.43 |
| FL_OFC_POG_r | 3.27 ± 0.38 | 1.20 ± 0.27 | | | |
| FL_strai_G_l | 2.69 ± 0.34 | 0.60 ± 0.13 | | | |
| FL_strai_G_r | 2.88 ± 0.34 | 0.77 ± 0.20 | | | |
| Subgen_antCing_l | 0.78 ± 0.13 | 0.59 ± 0.10 | | | |
| Subgen_antCing_r | 0.69 ± 0.13 | 0.54 ± 0.10 | | | |
| Subcall_area_l | 0.21 ± 0.04 | 0.02 ± 0.01 | | | |
| Subcall_area_r | 0.18 ± 0.03 | 0.02 ± 0.01 | | | |
| Presubgen_antCing_l | 0.76 ± 0.11 | 0.11 ± 0.05 | | | |
| Presubgen_antCing_r | 0.50 ± 0.07 | 0.08 ± 0.05 | | | |

### Regional analysis

Figure 3 and Figure 4 show boxplots of mean regional SUV and SUVr respectively, extracted in a selection of grey matter anatomical regions, for all subjects in the database. Each region is composed of left and right sub-regions. Mean regional SUV values were 5.36 ± 1.32, range 1.35 - 8.54 (Figure 3). Three subjects in the database had lower SUV values (between 1.35 to 3).

**Figure 3:**
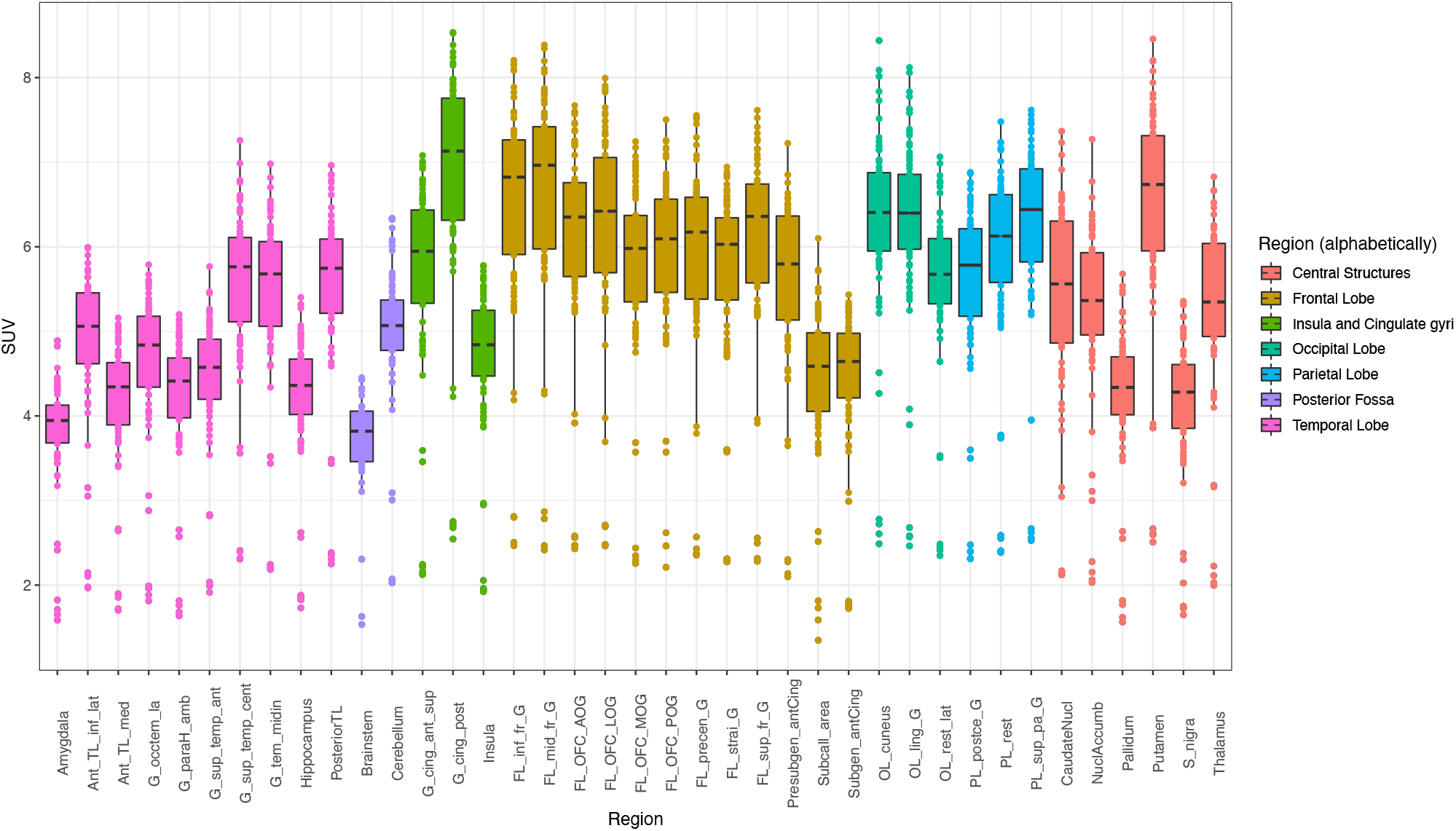
Boxplot of regional SUV for all subjects in the database. Centre lines correspond to medians, boxes to interquartile ranges, and whiskers to robust ranges. Outliers are represented as dots. Each dot represents a participant for unpaired regions and a participant’s right or left SUV value for paired regions.

**Figure 4:**
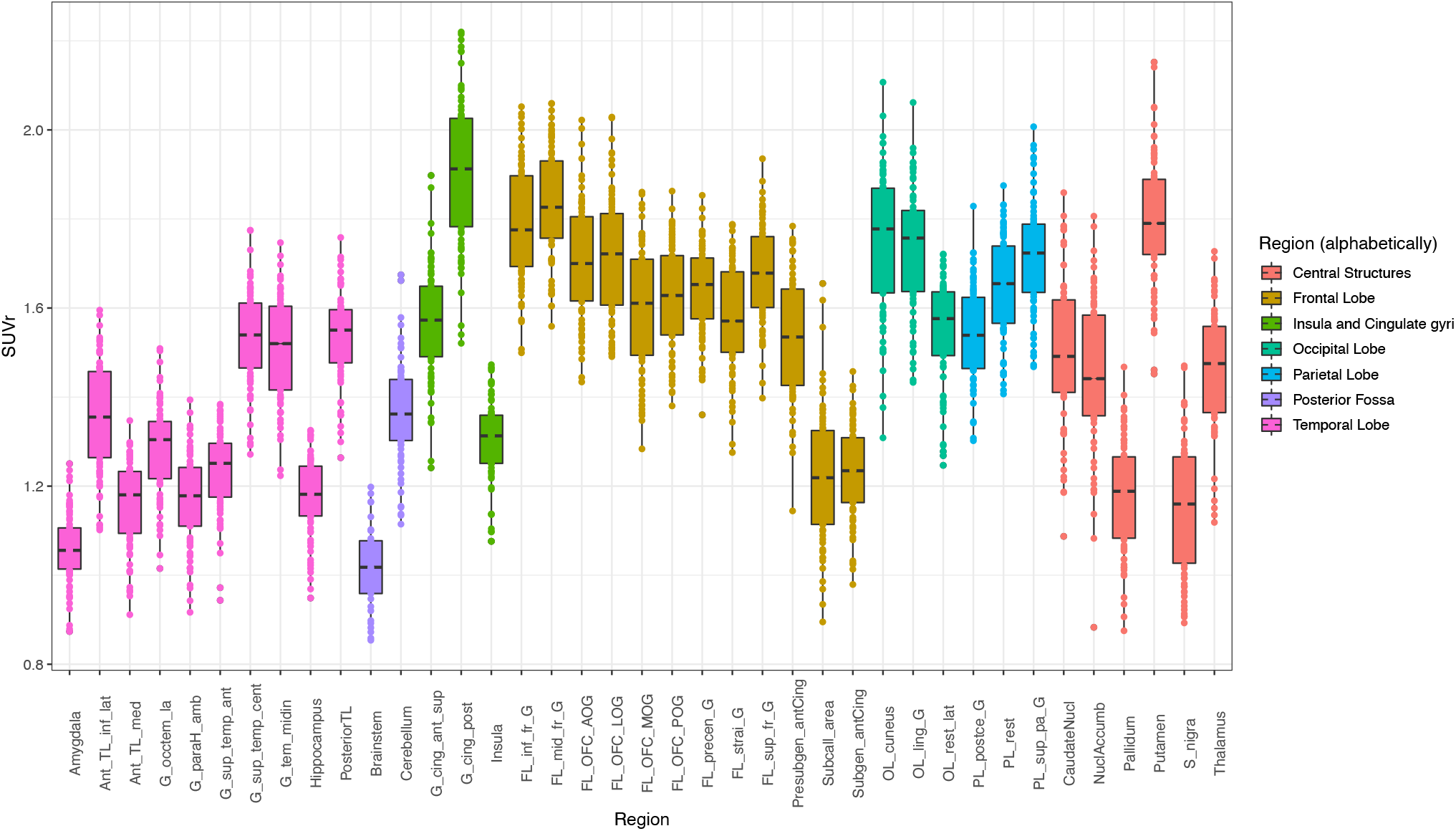
Boxplot pf regional activity normalized by mean activity in ICV mask (SPM) for all subjects in the database. Centre lines correspond to medians, boxes to interquartile ranges, and whiskers to robust ranges. Outliers are represented as dots.

The distribution of SUVr values (Figure 4) remains very similar to the distribution of SUV values (1.49 mean ± 0.26 SD, range 0.85 - 2.22), except that the dispersion is reduced and the outlier values from the three participants with unusually low SUVs are regularized.

Normalizing with the ICV mean value thus acts as an efficient way for regularizing the SUV distribution leaving the inter-regional variability intact.

### Leave-one-out SPM analysis

Results for the leave-one-out analysis of [^18^F]FDG PET are reported in Table 2.

**Table 2:** % of abnormality in the database. Cluster-level and voxel-level results are reported at p < 0.05 FWE. The denominator for subject-level is the total number of participants; the denominator for the cluster-level is the average number of resolution elements in the mask; the denominator for the voxel-level is the number of voxels in the SPM mask. See Methods for details.

| Contrast | Subject-level | Cluster-level | Voxel-level |
| --- | --- | --- | --- |
| **Hypermetabolism** | 13.5% (n=5) | 0.93% | 0.03% |
| **Hypometabolism** | 29.7% (n=11) | 5.21% | 0.32% |

At the subject-level, 5/37 (13.5%) of the participants had any significant increases in [^18^F]FDG uptake (hypermetabolism) relative to the other 36 participants. Any significant decreases (hypometabolism) was found for 11/37 (29.7%) of the participants.

At the cluster-level, significant changes were found in at most 5.21% of resolution elements, and at the voxel-level, in at most 0.32% of voxels.

All abnormalities in controls compared with controls are by definition false positives. We examined all 16 and present our findings in the Appendix (Table A1 and Table A2). Virtually all false positives had an anatomical or artefactual explanation.

## Discussion

A new database of 37 healthy subjects including T1 and FLAIR MRI, CT, and [^18^F]FDG PET images, called IDB-MRXFDG, has been created.

The age range has been selected to reflect the ages of participants in cognitive and clinical research studies at the CERMEP imaging centre, encompassing amongst others epilepsy, movement disorders, multiple sclerosis and disorders of consciousness and will align with the research priorities of many similar centres.

We performed quality control of all images visually and by screening for volumetric and regional SUV abnormalities. Three subjects had unusually low SUVs; this may be due to imperfect observation of the need for fasting ahead of the scan. We show that a simple global normalisation procedure removes the resulting outliers (Figure 4); depending on the application more sophisticated intra-scan normalisation procedures are conceivable (Yakushev et al. 2009, 2008). We also performed SPM leave-one-out studies for [^18^F]FDG. The relatively high false-positive rates per subject are explained by the existence of significant clusters of small size (from 1 to 95 voxels). Areas of apparent hypermetabolism were either at the edge of the brain or at the bottom of a particularly deep sulcus (see appendix, Table A1); areas of apparent hypometabolism (Table A2) were clearly linked to the participant’s anatomy, typically to a wide sulcus or fissure (7/11 cases). The other 4 cases were extracerebral or at the edge of the brain, probably linked to imperfect normalisation. We believe none would have been considered abnormal had they been seen in an analysis comparing one research subject with a particular condition against a group of controls. When testing the normality of the database at the cluster and voxel-level, the expected threshold of 5% of abnormality or lower was found for both hyper- and hypo-metabolism. The database therefore appears suitable for voxel-based [^18^F]FDG PET analysis with a ≤5% risk of Type 1 error.

The IDB-MRXFDG database could be used in many different applications such as the statistical comparison of a patient (or group of patients) to a database of healthy subjects, automatic quantitative analyses, and more generally methodology development in neuroimaging.

The inclusion of [^18^F]FDG PET in IDB-MRXFDG is particularly important. While there are now many MR databases covering, with varying density, the human lifespan as reviewed in (Coupé et al. 2017), we are aware of very few [^18^F]FDG PET databases. Wei et al. (Wei et al. 2018) scanned 78 healthy subjects aged 3-78 years on a PET/CT scanner; it is not clear whether this database is available on request, and there is no mention of MRI. The Marseille database (used e.g. in (Eusebio et al. 2012)) contains data from 60 healthy adults aged 21-78; [^18^F]FDG PET, T1 weighted MRI, and CT data are available by arrangement. A rare paediatric database (Archambaud et al. 2013) contains 24 datasets of participants aged 4.5-17.9 years (mean ± SD 10.06 ±3.1 years) and may be shared on request. These are “pseudocontrols” derived from epilepsy patients, selected from among a total of 71 children as the subgroup with both a normal visual analysis and a normal SPM analysis derived iteratively. They have been scanned on a traditional PET scanner with transmission-based attenuation correction which makes comparison with PET/CT data difficult (Sousa et al. 2020); no MRI is available. A large database available on request is the Alzheimer’s Disease Neuroimaging Initiative, ADNI (http://adni.loni.usc.edu/about/) which comprises over 300 healthy control [^18^F]FDG PET datasets; however, participants are aged 55-90 and therefore more suited to dementia research but outside the typical age range used for studies in normal cognition or epilepsy, one of the main clinical indications for clinical brain FDG PET. Similar concerns about the age of participants apply to those databases from the world-wide ADNI (WW-ADNI) networks that do contain FDG, as for example the Japan ADNI (J-ADNI; age 60-84) (Iwatsubo et al. 2018).

Examples of database uses for work in MR include the *voxel-wise* comparison of a patient with a control group to detect abnormalities from T1 images via voxel-based morphometry (Ashburner and Friston 2000; Richardson et al. 1997) and its variants that use T1 derivatives like grey-white matter junction images (Antel et al. 2002; Huppertz et al. 2005) for the detection of specific pathologies like Focal Cortical Dysplasia. FLAIR as a sequence highly sensitive to pathology has similarly been used at the single-subject level in comparison to control groups (e.g. (Focke et al. 2008; Huppertz et al. 2011)). Another group of examples is the *region-wise* comparison of the size of cerebral structures between groups or between individuals and a control group (e.g. (Heckemann et al. 2011; Hammers et al. 2007; Sapey-Triomphe et al. 2015; Klein-Koerkamp et al. 2014)). Importantly, such work has been successfully undertaken with control groups scanned on a different scanner (e.g. (Cross et al. 2013; Yaakub et al. 2020)), and IDB-MRXFDG could be used to increase the size of control groups.

Since PET-CT scanners rapidly displaced PET-only scanners in the early 2000s, low-dose CT has been coupled to brain [^18^F]FDG PET for estimation of tissue density and attenuation correction. With the advent of commercial PET-MR scanners since 2011, there has been no direct way of measuring electron density in the head, and alternative approaches have had to be found. Synthesis of “pseudo-CTs” via atlas approaches (Burgos et al. 2014; Merida et al. 2017) is a successful approach that performs well overall (Ladefoged et al. 2017) but requires pairs of MR and CT images to achieve the synthesis. IDB-MRXFDG has already been used for such approaches (Merida et al. 2015).

The latter application of databases – MR-based attenuation based on MR-CT pairs – is one domain where Deep Learning methods, notably with Convolutional Neuronal Networks, have recently become very successful (Ladefoged et al. 2020; Yaakub, McGinnity, Beck, et al. 2019). However, they often require substantially larger training datasets or priors than multi-atlas methods, in the case of MR-based attenuation recently estimated at 100-400 pairs, with an influence of MR heterogeneity (Ladefoged et al. 2020). More widespread availability of databases will further Deep Learning approaches, particularly when multiple modalities are available per subject, allowing e.g. synthesis of missing modalities (Yaakub, McGinnity, Clough, et al. 2019).

## Data sharing

We have obtained Ethical permission to make the database available on request for *bona fide* research. Please email for a short access form, detailing which format you require (DICOM format, NIFTI in subject’s space and NIFIT in normalized space, following BIDS common specification). Your request will then be considered by the access committee. If found in line with permitted use (i.e. *bona fide* research) a licence will be issued and the requested database transferred securely.

## Appendix

**Table A1:**
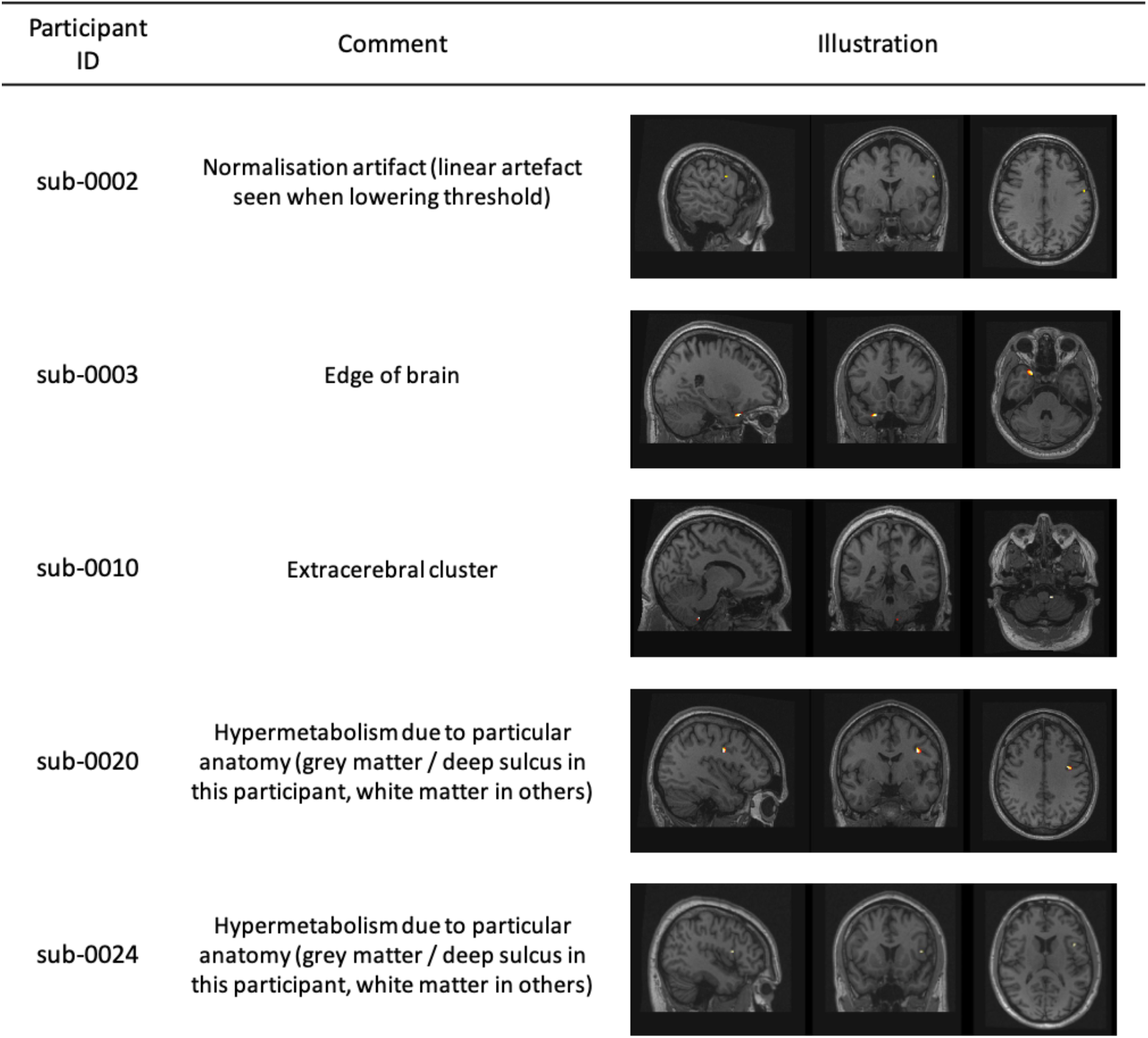
Individual thresholded T-maps (p < 0.05 FWE) for participants showing significant **increases** in [^18^F]FDG uptake (hypermetabolism) relative to the other 36 participants (false positives). The analysis consisted in a leave-one-out ANCOVA performed on SPM12 (see Methods for details). For each case, we provide an anatomical or artefactual explanation.

| Participant ID | Comment | Illustration |
| --- | --- | --- |
| sub-0002 | Normalisation artifact (linear artefact seen when lowering threshold) |  |
| sub-0003 | Edge of brain |  |
| sub-0010 | Extracerebral cluster |  |
| sub-0020 | Hypermetabolism due to particular anatomy (grey matter / deep sulcus in this participant, white matter in others) |  |
| sub-0024 | Hypermetabolism due to particular anatomy (grey matter / deep sulcus in this participant, white matter in others) |  |

**Table A2:**
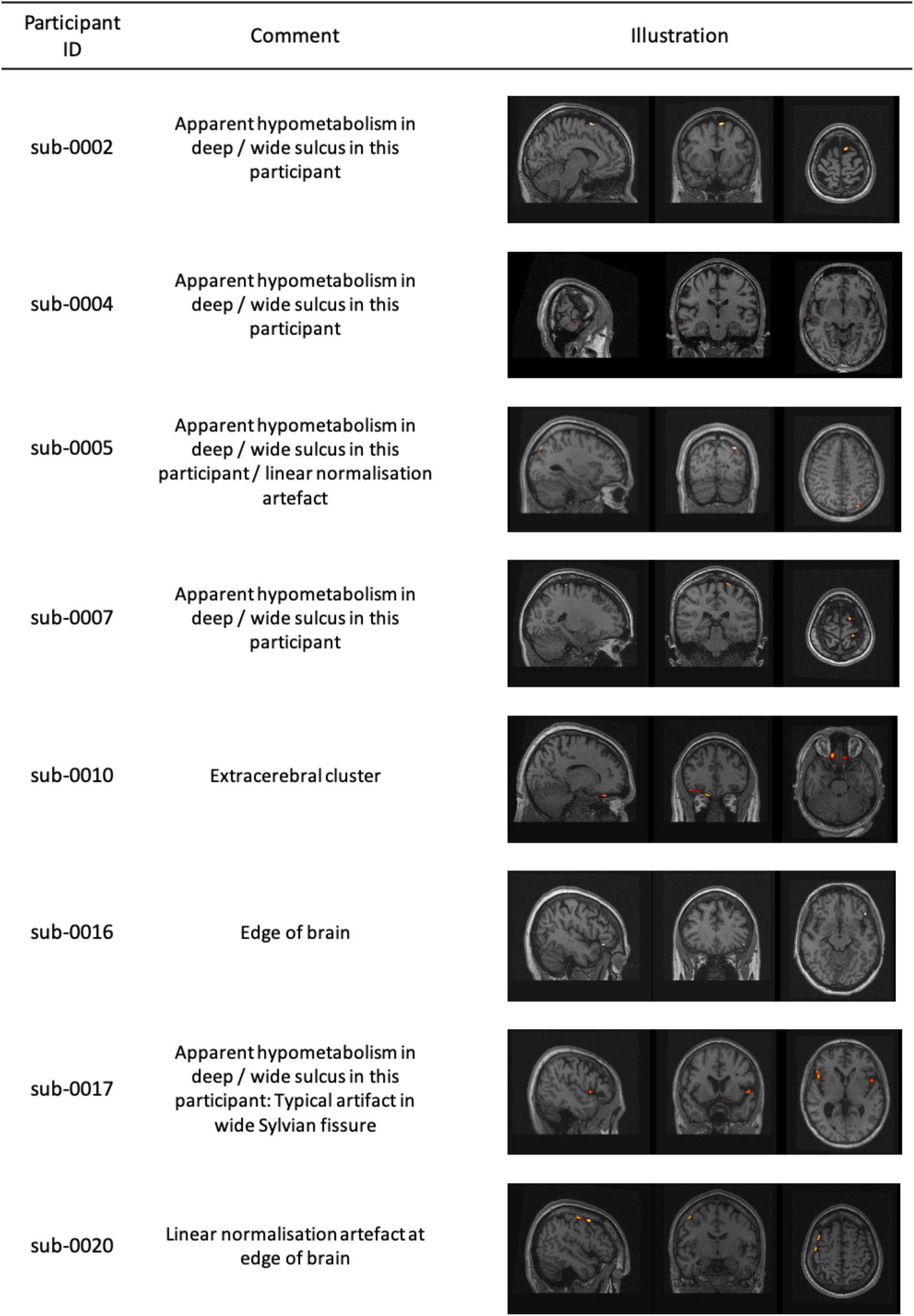
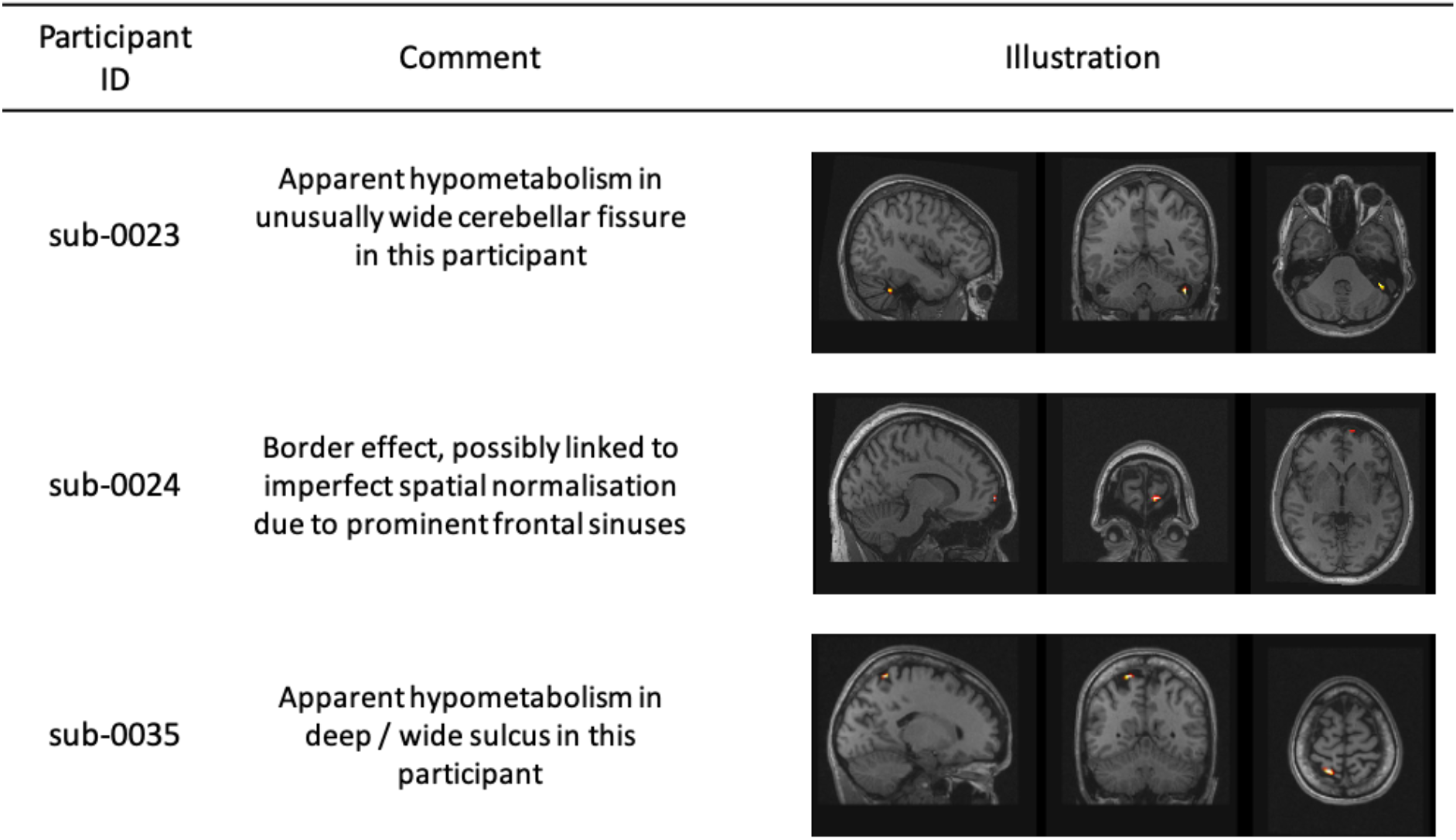
Individual thresholded T-maps (p < 0.05 FWE) for participants showing significant **decreases** in [^18^F]FDG uptake (hypometabolism) relative to the other 36 participants (false positives). The analysis consisted in a leave-one-out ANCOVA performed on SPM12 (see Methods for details). For each case, we provide an anatomical or artefactual explanation.

| Participant ID | Comment | Illustration |
| --- | --- | --- |
| sub-0002 | Apparent hypometabolism in deep / wide sulcus in this participant |  |
| sub-0004 | Apparent hypometabolism in deep / wide sulcus in this participant |  |
| sub-0005 | Apparent hypometabolism in deep / wide sulcus in this participant / linear normalisation artefact |  |
| sub-0007 | Apparent hypometabolism in deep / wide sulcus in this participant |  |
| sub-0010 | Extracerebral cluster |  |
| sub-0016 | Edge of brain |  |
| sub-0017 | Apparent hypometabolism in deep / wide sulcus in this participant: Typical artifact in wide Sylvian fissure |  |
| sub-0020 | Linear normalisation artefact at edge of brain |  |
| sub-0023 | Apparent hypometabolism in unusually wide cerebellar fissure in this participant |  |
| sub-0024 | Border effect, possibly linked to imperfect spatial normalisation due to prominent frontal sinuses |  |
| sub-0035 | Apparent hypometabolism in deep / wide sulcus in this participant |  |

**Table A3:** Abbreviation list of the 83 regions used in the ROI evaluation based on the Hammers_mith atlases (www.brain-development.org/ Hammers et al. 2003, Gousias et al. 2008)

| Abbreviation | Complete name |
| --- | --- |
| Hippocampus_r | Hippocampus (right) |
| Hippocampus_l | Hippocampus (left) |
| Amygdala_r | Amygdala (right) |
| Amygdala_l | Amygdala (left) |
| Ant_TL_med_r | Anterior temporal lobe, medial part (right) |
| Ant_TL_med_l | Anterior temporal lobe, medial part (left) |
| Ant_TL_inf_lat_r | Anterior temporal lobe, lateral part excluding superior temporal gyrus (right) |
| Ant_TL_inf_lat_l | Anterior temporal lobe, lateral part excluding superior temporal gyrus (left) |
| G_paraH_amb_r | Parahippocampal and ambient gyri (right) |
| G_paraH_amb_l | Parahippocampal and ambient gyri (left) |
| G_s_t_cent_r | Superior temporal gyrus, central part (right) |
| G_s_t_cent_l | Superior temporal gyrus, central part (left) |
| G_tem_midin_r | Middle and inferior temporal gyrus (right) |
| G_tem_midin_l | Middle and inferior temporal gyrus (left) |
| G_occtem_la_r | Fusiform (lateral occipitotemporal) gyrus (right) |
| G_occtem_la_l | Fusiform (lateral occipitotemporal) gyrus (left) |
| Cerebellum_r | Cerebellum (right) |
| Cerebellum_l | Cerebellum (left) |
| Brainstem | Brainstem |
| Insula_l | Insula (left) |
| Insula_r | Insula (right) |
| OL_rest_lat_l | Lateral remainder of occipital lobe (left) |
| OL_rest_lat_r | Lateral remainder of occipital lobe (right) |
| G_cing_a_s_l | Cingulate gyrus, anterior part (left) |
| G_cing_a_s_r | Cingulate gyrus, anterior part (right) |
| G_cing_p_l | Gyrus cinguli, posterior part (left) |
| G_cing_p_r | Gyrus cinguli, posterior part (right) |
| FL_mid_fr_G_l | Middle frontal gyrus (left) |
| FL_mid_fr_G_r | Middle frontal gyrus (right) |
| PosteriorTL_l | Posterior temporal lobe (left) |
| PosteriorTL_r | Posterior temporal lobe (right) |
| PL_rest_l | Inferiolateral remainder of parietal lobe (left) |
| PL_rest_r | Inferiolateral remainder of parietal lobe (right) |
| CaudateNucl_l | Caudate nucleus (left) |
| CaudateNucl_r | Caudate nucleus (right) |
| NuclAccumb_l | Nucleus accumbens (left) |
| NuclAccumb_r | Nucleus accumbens (right) |
| Putamen_l | Putamen (left) |
| Putamen_r | Putamen (right) |
| Thalamus_l | Thalamus (left) |
| Thalamus_r | Thalamus (right) |
| Pallidum_l | Pallidum (left) |
| Pallidum_r | Pallidum (right) |
| Corp_Callosum | Corpus callosum |
| LatVent_excl_TH_r | Lateral ventricle (excluding temporal horn) (right) |
| LatVent_excl_TH_l | Lateral ventricle (excluding temporal horn) (left) |
| BodyVentricle_r | Lateral ventricle, body of right ventricle |
| BodyVentricle_l | Lateral ventricle, body of left ventricle |
| ThirdVentricle | Third ventricle |
| FL_precen_G_l | Precentral gyrus (left) |
| FL_precen_G_r | Precentral gyrus (right) |
| FL_strai_G_l | Straight gyrus (left) |
| FL_strai_G_r | Straight gyrus (right) |
| FL_OFC_AOG_l | Anterior orbital gyrus (left) |
| FL_OFC_AOG_r | Anterior orbital gyrus (right) |
| FL_i_fr_G_l | Inferior frontal gyrus (left) |
| FL_i_fr_G_r | Inferior frontal gyrus (right) |
| FL_s_fr_G_l | Superior frontal gyrus (left) |
| FL_s_fr_G_r | Superior frontal gyrus (right) |
| PL_postce_G_l | Postcentral gyrus (left) |
| PL_postce_G_r | Postcentral gyrus (right) |
| PL_s_pa_G_l | Superior parietal gyrus (left) |
| PL_s_pa_G_r | Superior parietal gyrus (right) |
| OL_ling_G_l | Lingual gyrus (left) |
| OL_ling_G_r | Lingual gyrus (right) |
| OL_cuneus_l | Cuneus (left) |
| OL_cuneus_r | Cuneus (right) |
| FL_OFC_MOG_l | Medial orbital gyrus (left) |
| FL_OFC_MOG_r | Medial orbital gyrus (right) |
| FL_OFC_LOG_l | Lateral orbital gyrus (left) |
| FL_OFC_LOG_r | Lateral orbital gyrus (right) |
| FL_OFC_POG_l | Posterior orbital gyrus (left) |
| FL_OFC_POG_r | Posterior orbital gyrus (right) |
| S_nigra_l | Substantia nigra (left) |
| S_nigra_r | Substantia nigra (right) |
| Subgen_antCing_l | Subgenual frontal cortex (left) |
| Subgen_antCing_r | Subgenual frontal cortex (right) |
| Subcall_area_l | Subcallosal area (left) |
| Subcall_area_r | Subcallosal area (right) |
| Presubgen_antCing_l | Pre-subgenual frontal cortex (left) |
| Presubgen_antCing_r | Pre-subgenual frontal cortex (right) |

## Notes

### Competing Interest Statement

The authors have declared no competing interest.

